# Insights into the role of glycerophospholipids on the iron export function of SLC40A1 and the molecular mechanisms of ferroportin disease

**DOI:** 10.1101/2024.02.05.578958

**Authors:** Rim Debbiche, Ahmad Elbahnsi, Kévin Uguen, Chandran Ka, Isabelle Callebaut, Gérald Le Gac

**Author notes:** **Corresponding Author:** Gerald Le Gac, Institut Brestois de Recherche en Bio-Santé, UFR Médecine et Sciences de la Santé, UMR1078, 22 rue Camille Desmoulins, 29238 Brest, France. Inserm U1268 MCTR, CiTCoM UMR 8038 CNRS, Université Paris Cité, 75006 Paris, France. these authors contribute equally (Rim Debbiche, Ahmad Elbahnsi, Isabelle Callebaut, Gérald Le Gac). **Authorship contributions** R. Debbiche, I. Callebaut and G. Le Gac, designed the study. R. Debbiche, A. Elbahnsi and I. Callebaut conducted experiments. R. Debbiche, A. Elbahnsi, K. Uguen, C. Ka, I. Callebaut and G. Le Gac analyzed data. R. Debbiche, A. Elbahnsi, I. Callebaut and G. Le Gac, wrote the manuscript. All authors contributed to the editing of the final manuscript.

## Abstract

SLC40A1 is the sole iron export protein reported in mammals. In humans, its dysfunction is responsible for ferroportin disease, an inborn error of iron metabolism transmitted as an autosomal dominant trait and observed in different ethnic groups. As a member of the major facilitator superfamily, SLC40A1 requires a series of conformational changes to enable iron translocation across the plasma membrane. The influence of lipids on these conformational changes has been little investigated to date. Here, we combine molecular dynamics simulations of SLC40A1 embedded in bilayers with experimental alanine scanning mutagenesis to analyze the specific role of glycerophospholipids. We identify four basic residues (Lys90, Arg365, Lys366 and Arg371) that are located at the membrane-cytosol interface and consistently interact with POPC and POPE lipid molecules. These residues surround a network of salt bridges and hydrogens bonds that play a critical role in stabilizing SLC40A1 in its basal outward-facing conformation. More deeply embedded in the plasma membrane, we identify Arg179 as a charged amino acid residue also tightly interacting with lipid phosphates. This result into a local deformation of the lipid bilayer. Interestingly, Arg179 is adjacent to Arg178, which forms a functionally important salt-bridge with Asp473 and is a recurrently associated with ferroportin disease when mutated to glutamine. We demonstrate that the two p.Arg178Gln and p.Arg179Thr missense variants have similar functional behaviors. These observations provide insights into the role of phospholipids in the formation/disruption of the SLC40A1 inner gate, and give a better understanding of the diversity of molecular mechanisms of ferroportin disease.

## INTRODUCTION

Ferroportin 1, also referred to as Solute Carrier Family 40 Member 1 (SLC40A1), is essential for proper maintenance of human iron homeostasis at systemic and cellular levels. This 62,5 kDa multipass plasma membrane protein, which is notably expressed in reticuloendothelial macrophages, duodenal enterocytes, hepatocytes, placenta syncytiotrophoblasts and erythrocytes, is the only mammalian iron exporter (1–5). Its dysfunction is responsible for Ferroportin Disease (FD), an inborn error of iron metabolism transmitted as an autosomal dominant trait and observed in different ethnic groups (6–8).

SLC40A1 adopts the canonical Major Facilitator Superfamily (MFS)-fold that consist in 12 transmembrane (TM) helices, organized into two structurally similar bundles of 6 TMs each, which exhibit pseudo-symmetry. Each bundle is made up from 3 TM structural inverted repeats (N-terminal domain: TMs 1-3 and TMs 4-6; C-terminal domain: TMs 7-9 and TMs 10-12) (9, 10), with a specific role for each TM: TM 1,4,7,10 are the core helices, TM 2,5,8,11 the interfacial helices, and TM 3,6,9,12 the external helices. In the basal state, the two 6-TMs bundles of SLC40A1 form an outward-facing (OF) open conformation, with TM4 and TM10 at the center of the closed, inner gate. SLC40A1 undergoes large conformational changes to take up cytosolic iron and deliver it into the extracellular milieu. According to the MFS “clamp-and-switch” transport cycle model, these conformational changes involve both rigid-body rotations and local rearrangements. The conformational transitions are orchestrated by the formation and disruption of non-covalent interactions between the N- and C-terminal domain, allowing for a succession of (at least) five conformational states: outward-facing, outward-occluded, occluded, inward-occluded, inward-facing (9, 11, 12).

Based on a comparative modelling strategy and molecular dynamics simulations, we previously highlighted that motif A, which is the best-conserved sequence element of the MFS family, is the critical point of two networks of salt bridges and hydrogen bonds that link together the N- and C-domains in the inner leaflet of the plasma membrane and play both structural and functional roles in stabilizing human SLC40A1 in its OF conformation (13). BillesbØlle *et al*. reported the first cryo-electron microscopy 3D structures of human SLC40A1 in an OF conformation embedded in lipid nanodiscs (14), followed by that of the primate Philippine Tarsier in the same conditions (15). The authors confirmed the critical role of intra- and inter-lobe ion-pair interactions for inner gate stability, and a consistent involvement in conformational transitions between different states. We also pointed out the importance of gating residues in understanding the molecular mechanisms of FD (13), with a particular focus on the p.Arg178Gln missense mutation which breaks an important inter-domain salt bridge between Arg178 (TM5) and Asp157 (TM10) and is probably the most common disease-causing mutation reported to date (8, 16).

Questions still remain unanswered about the molecular mechanisms associated with iron translocation across the plasma membrane. One of the most important is how SLC40A1 manages to overcome energy barriers within the lipid bilayer. A first hypothesis is that SLC40A1 utilizes the so-called “proton-motive force”, as many other MFS proteins (15, 17, 18). A second hypothesis, which we have formulated earlier (13), is that non-covalent interactions between polar and charged amino acids act as molecular switches that drive a series of conformational transitions between the two extremes OF and IF conformations, and that these charge/polar clusters are regulated by protein-lipid interactions (13).

The present study aimed at identifying charged residues in the vicinity of the human SLC40A1 inner gate that could interact directly with glycerophospholipids and be involved in conformational transitions. Based on molecular dynamics (MD) simulations performed on the experimental 3D structure of human SLC40A1 in the OF conformation, we provide initial clues by focusing on five basic amino acids (Lys90, Arg179, Arg365, Lys366 and Arg371) and undertaking functional studies. Interestingly, the p.Arg179Thr substitution has been reported in an Italian patient with unexplained hyperferritinemia (19). We investigate this variant and provide a more definitive demonstration of its involvement in FD.

## METHODS

### SLC40A1 plasmid constructs

The wild-type (WT) FPN1-V5 and FPN1-V5/CD8 bicistronic plasmid constructs were generated as previously described (13, 16). All FPN1 mutations were introduced in the different vectors by using the QuikChange Site-Directed mutagenesis kit, according to the manufacturer’s instructions (Agilent Technologies). Sequencing analyses were performed to check the integrity of all plasmid constructs (full length *SLC40A1* cDNA sequenced after each site-directed mutagenesis).

### Culture and transfection of human epithelial kidney (HEK)293T cells

Human epithelial kidney (HEK) 293T cells, from the American Type Culture Collection, were incubated at 37°C in a 5% CO2 humidified atmosphere and propagated in Dulbecco’s modified Eagle’s medium (DMEM; Lonza) supplemented with 10% fetal bovine serum. Cells were transiently transfected using Lipofectamine 2000 (ThermoFisher Scientific; flow cytometry experiments) or Transit-2020 (Mirus Bio LLC; iron export experiments), according to the manufacturer’s instructions, and a 3:1 transfection reagent (µL)/plasmid DNA ratio (µg).

### Flow cytometry analysis

Flow cytometry experiments were done as previously described (13). Briefly, HEK293T cells (1.75 × 10^5^ cells/well in 6-well plates) transfected with the pIRES_FPN1-V5_CD8 constructs were treated (or not) with 4.3 nM of native human-25 hepcidin (24h after transfection) for 16 hours. Cells were harvested with trypsin, pelleted (500 g, 5 min, 4°C) and resuspended in PBS (pH 7.4) containing EDTA (Lonza) and 10% fetal bovine serum, before being incubated for 20 min at 4°C with anti-V5-FITC or anti-V5-PE (ThermoFisher Scientific) and anti-CD8-APC (Miltenyi Biotec). Stained cells were pelleted (500 g, 5 min, 4°C) and resuspended in 400 *µ*L PBS-EDTA. Cells were analyzed using a BD Accuri C6 flow cytometer (BD Biosciences) and FlowLogic™ software (Miltenyi Biotec).

### ^55^Fe release measurements

^55^Fe loading of human apotransferrin was performed as previously described (20). Briefly, HEK293T cells (1.7 × 10^5^ cells/well in 12-well plates) were grown for 24 h in supplemented DMEM (Lonza), before to be incubated with 20 µg/mL ^55^Fe-transferrin for 24 h and transiently transfected with wild-type or mutated FPN1-V5 plasmid constructs. Fifteen hours post-transfection cells were washed once with PBS and cultured in Pro293a-CDM serum-free medium (BioWhittaker) for up to 36 h. ^55^Fe exported into the supernatant was collected, mixed with liquid scintillation fluid (Ultima Gold MV, Packard Bioscience) and counted for 10min in a TRICARB 1600 CA scintillation counter (Packard). Percentage of ^55^Fe export was calculated using the following formula: (^55^Fe in the supernatant at end point, divided by cellular ^55^Fe at time zero) x 100.

### Molecular dynamics simulation and 3D structure analysis

Molecular dynamics (MD) simulations were carried out using the experimental 3D structure of human FPN1 in its apo OF-state obtained by cryo-EM (pdb 6W4S). The wild type protein was prepared using the CHARMM-GUI server (21). The models were embedded in one of two types of lipid bilayers: either a mixture of POPC (1-palmitoyl-2-oleoyl-sn-glycero-3-phosphocholine), POPE (1-palmitoyl-2-oleoyl-sn-glycero-3-phosphoethanolamine), and cholesterol (CHOL) in a 2:1:1 ratio, or a simplified bilayer consisting only of POPC. The POPC:POPE:CHOL mixture systems were solvated using a solution of 150 mM NaCl + CaCl2, while the POPC-only systems were solvated in a 150 mM NaCl solution. The CHARMM36 force field was used for the protein, lipids and ions, and the TIP3P model for water (22). Minimization, equilibration and production steps were performed both on our local machines as well as on CINES supercomputers using different versions of Gromacs (2019, 2021 and 2023) (23). The standard CHARMM-GUI inputs were used for the minimization and equilibration of the systems. During these steps, harmonic restraints applied to the protein heavy atoms and the lipid heads and were gradually released during 1.2 ns. The production dynamics were then performed in the NPT ensemble without any restraints. Nose-Hoover thermostat and Parrinello-Rahman barostat to keep the temperature and the pressure constant at 310 K and 1 bar (24, 25). Periodic boundary conditions were used and the particle mesh ewald algorithm was applied to treat long-range electrostatic interactions (26). A switching function was applied between 10 and 12 Å for the non-bonded interactions. LINCS was applied to constrain the bond lengths involving hydrogen atoms (27). The integration timestep was set to 2 fs and the overall lengths of the trajectories were 500ns or 1 μs for the POPC:POPE:CHOL and POPC systems, respectively. Analysis of MD simulations, which included a total of four trajectories for the WT protein, was conducted by following the Root Mean Square Deviations (RMSD) of the C-alpha atoms. This calculation was performed for each simulation snapshot/frame in comparison to the initial cryo-EM conformation. Additionally, we measured the distances between critical residues using in-house tcl scripts. RMSD and distances figures were generated using R scripts.

3D structures were manipulated using Chimera 1.13.1. (28)and VMD.

### Statistical analysis

Data are presented as means (column bars) + standard deviation. Comparison used 2-tailed paired sample *t*-test.

## RESULTS

### MD-based prediction of lipid-interacting residues in human FPN1

Charged amino acids are not frequently observed in transmembrane regions. They can, however, contribute to the stability and dynamics of membrane proteins, forming specific non-covalent interactions with surrounding lipids. Interestingly, the frequency of positively charged amino acids (Arg, Lys, His) appears to be higher in the cytoplasmic part of eukaryotic plasma membrane proteins (29). In the present study, we focused on basic amino acids in the vicinity of the charge/polar clusters that form salt bridges and hydrogen bonds and favor assembly of the N- and C-domains in the OF basal state (highlighted in green in Figure 1).

**Figure 1.**
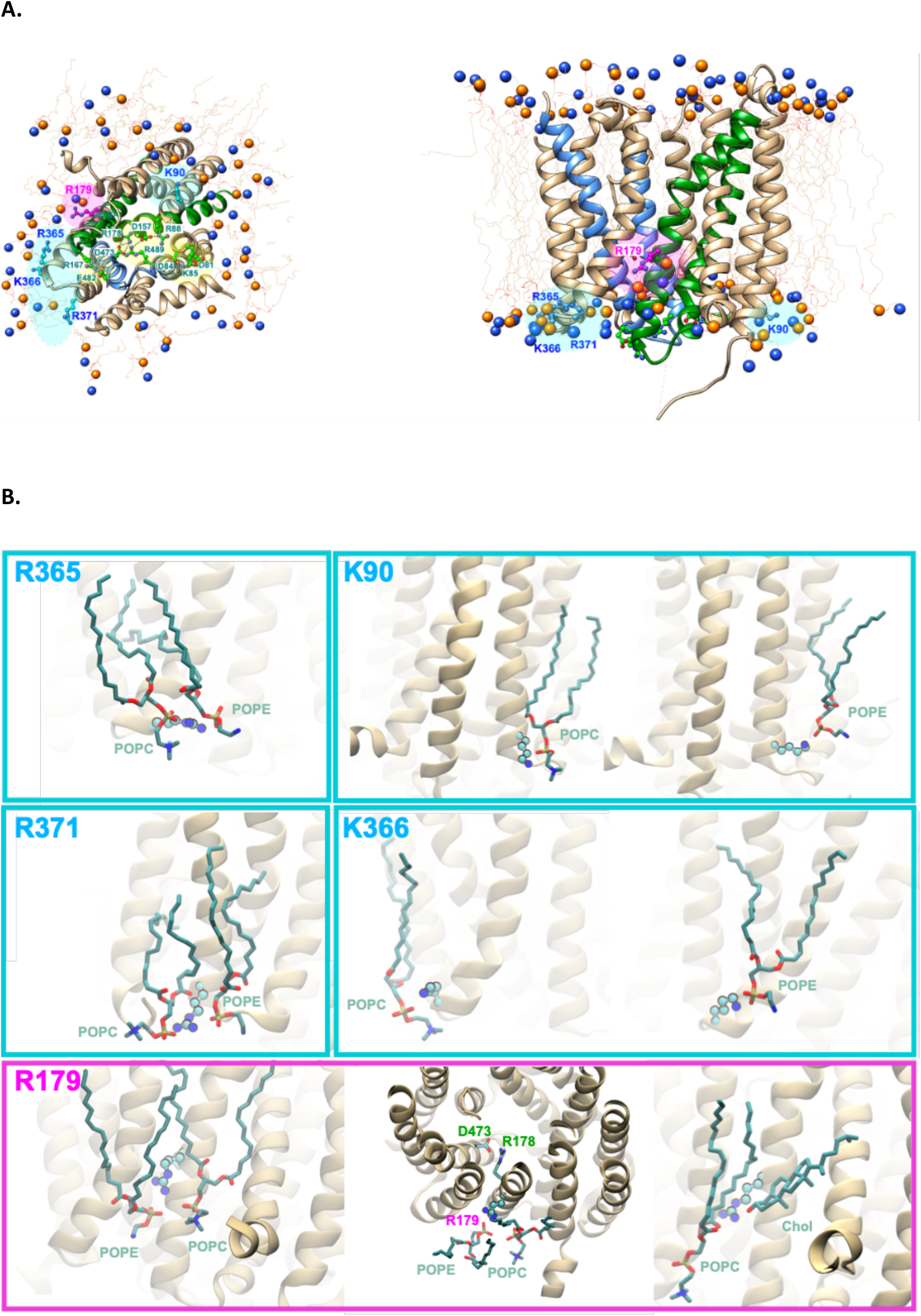
View of the 3D structure of human SLC40A1 embedded in a lipid bilayer. (A) Orthogonal views (ribbon representations) highlight the position of the five basic amino acids studied in this report (cyan and pink, ball-and-stick representation) on snapshots of the 3D structure of human SLC40A1 during 1µs MD simulation in presence of POPC, while amino acids of the inner gate are shown in green. TM4 and TM5 are colored in green, TM10 and TM11 are colored in blue. POPC molecules are represented in wireframe, with the position of their P and N atoms displayed as balls at left in order to visualize the localized bilayer deformation at the level of Arg179 (pink). (B) Details of the contacts observed in selected snapshots for each of the five amino acids (ball-and-stick representation), with the POPC and POPE headgroups (sticks) and with CHOL in the specific case of Arg179. The interaction of Arg179 with phospholipid headgroups result in their pulling towards the hydrophobic core of the bilayer. For convenience, amino acids are described by a 1-letter code, instead of the 3-letter code used in the main text.

We performed two 1µs-MD simulations using the experimental wild-type 3D structure of human SLC40A1 (pdb 6W4S) embedded in POPC lipid molecules. Two 500 ns-MD simulations were additionally conducted in a lipid bilayer composed of POPC, POPE and CHOL, in a 2:1:1 ratio, to better mimic the plasma membrane, as described in (30). These simulations provided statistical data for identifying possible interactions between positively charged amino acids and lipids. RMSD values for the C-alpha backbone indicate the overall stability of the system after equilibration (Supplementary Data 1).

Distances between the amino acids of the inner gate involved in critical salt-bridges (Arg178-Asp473; Arg88-Asp157-Arg489) show an overall stability along the simulation (Supplementary Data 2). Of note is the particular stability of the hydrogen bonds linking Asp174 and Gln478, which are located one helical turn below the critical salt-bridge between Arg178 and Asp473 (if SLC40A1 is viewed with its intracellular side down). Predicted positioning of lipids around SLC40A1 is, by contrast, highly dynamics. We have selected four positively charged amino acids (in blue in Figure 1), which are located at the interface between the lipid bilayer and the cytosol, within (Lys90-TM3; Ag371-TM9) or in the vicinity (Arg365 – TM8; Lys366 – TM8) of external helices. High-frequency contacts were observed for all four amino acids in the two POPC simulations (Table 1). Significant interactions with the lipid head groups of POPC and POPE where also detected in the MD simulations with a mixed lipid composition (Table 1 and snapshots in Figure 2B), whereas contacts with cholesterol were not, or only very rarely, observed.

**Table 1.**
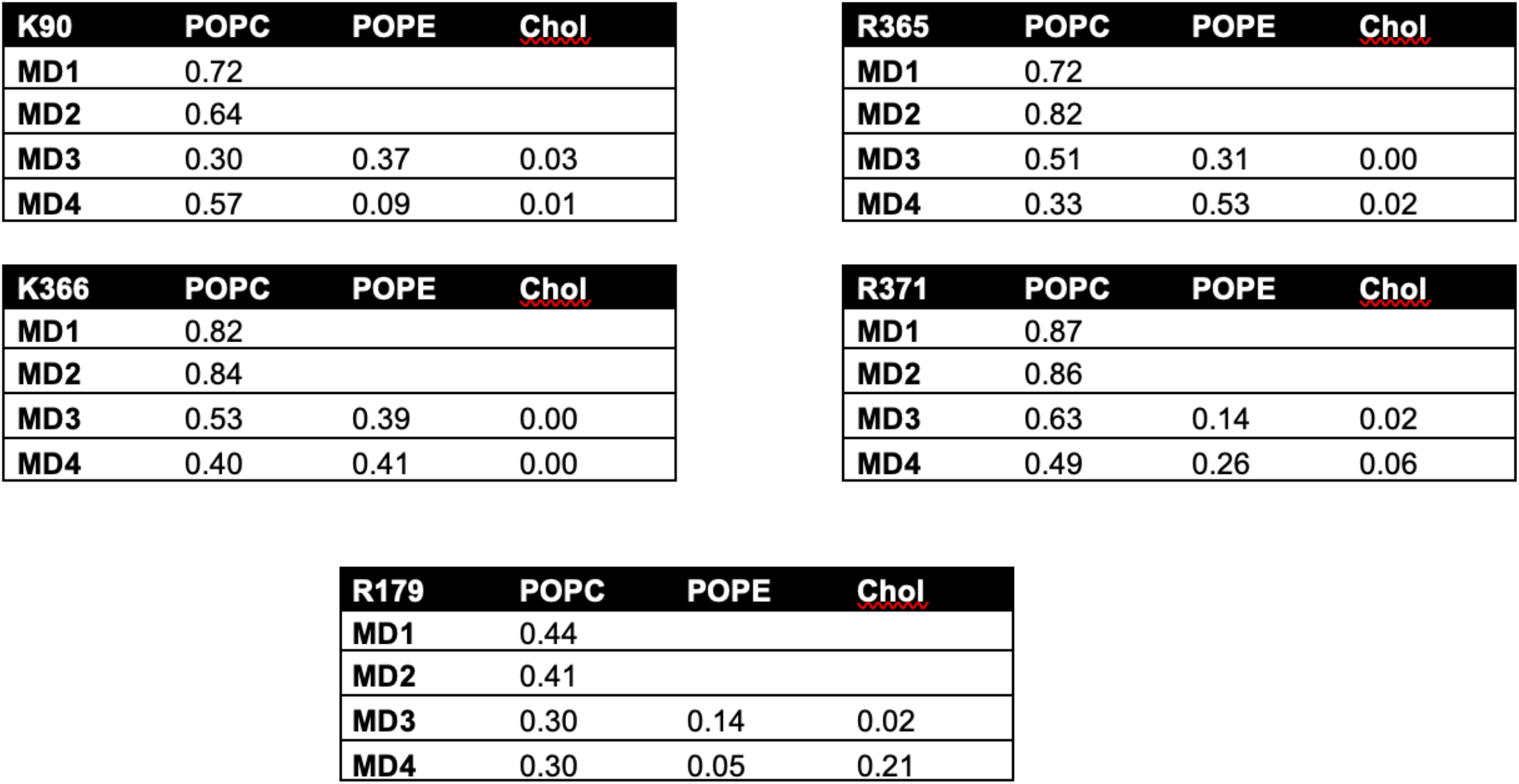
Contact frequency of the five basic amino acids with lipids. The frequencies of contacts between each basic amino acid and any lipid headgroups (P atoms), were determined by applying a distance cutoff of 5 Angstroms.

| **K90** | **POPC** | **POPE** | **Chol** |
| --- | --- | --- | --- |
| **MD1** | 0.72 |  |  |
| **MD2** | 0.64 |  |  |
| **MD3** | 0.30 | 0.37 | 0.03 |
| **MD4** | 0.57 | 0.09 | 0.01 |

| **R365** | **POPC** | **POPE** | **Chol** |
| --- | --- | --- | --- |
| **MD1** | 0.72 |  |  |
| **MD2** | 0.82 |  |  |
| **MD3** | 0.51 | 0.31 | 0.00 |
| **MD4** | 0.33 | 0.53 | 0.02 |

| **K366** | **POPC** | **POPE** | **Chol** |
| --- | --- | --- | --- |
| **MD1** | 0.82 |  |  |
| **MD2** | 0.84 |  |  |
| **MD3** | 0.53 | 0.39 | 0.00 |
| **MD4** | 0.40 | 0.41 | 0.00 |

| **R371** | **POPC** | **POPE** | **Chol** |
| --- | --- | --- | --- |
| **MD1** | 0.87 |  |  |
| **MD2** | 0.86 |  |  |
| **MD3** | 0.63 | 0.14 | 0.02 |
| **MD4** | 0.49 | 0.26 | 0.06 |

| **R179** | **POPC** | **POPE** | **Chol** |
| --- | --- | --- | --- |
| **MD1** | 0.44 |  |  |
| **MD2** | 0.41 |  |  |
| **MD3** | 0.30 | 0.14 | 0.02 |
| **MD4** | 0.30 | 0.05 | 0.21 |

**Figure 2-3.**
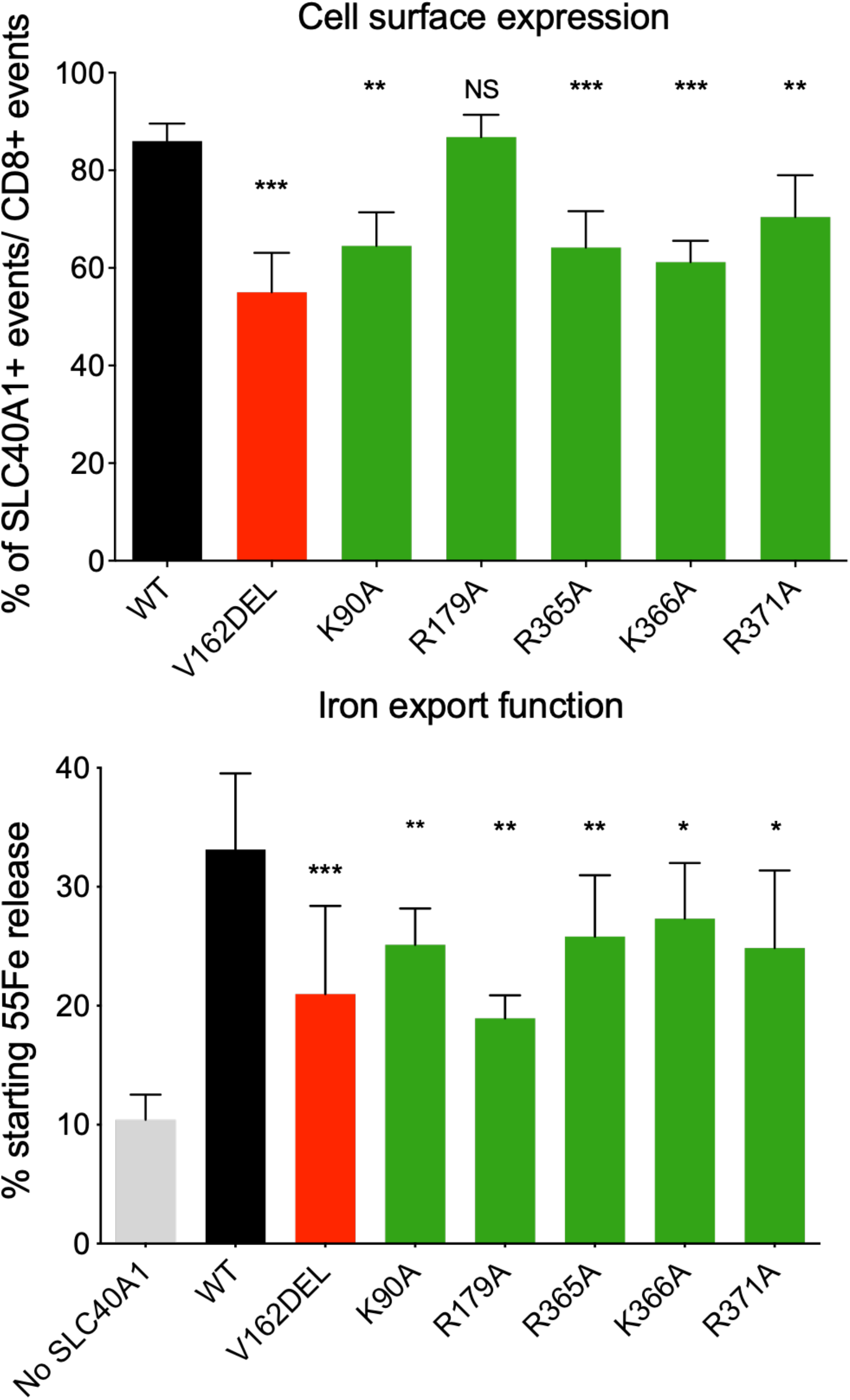
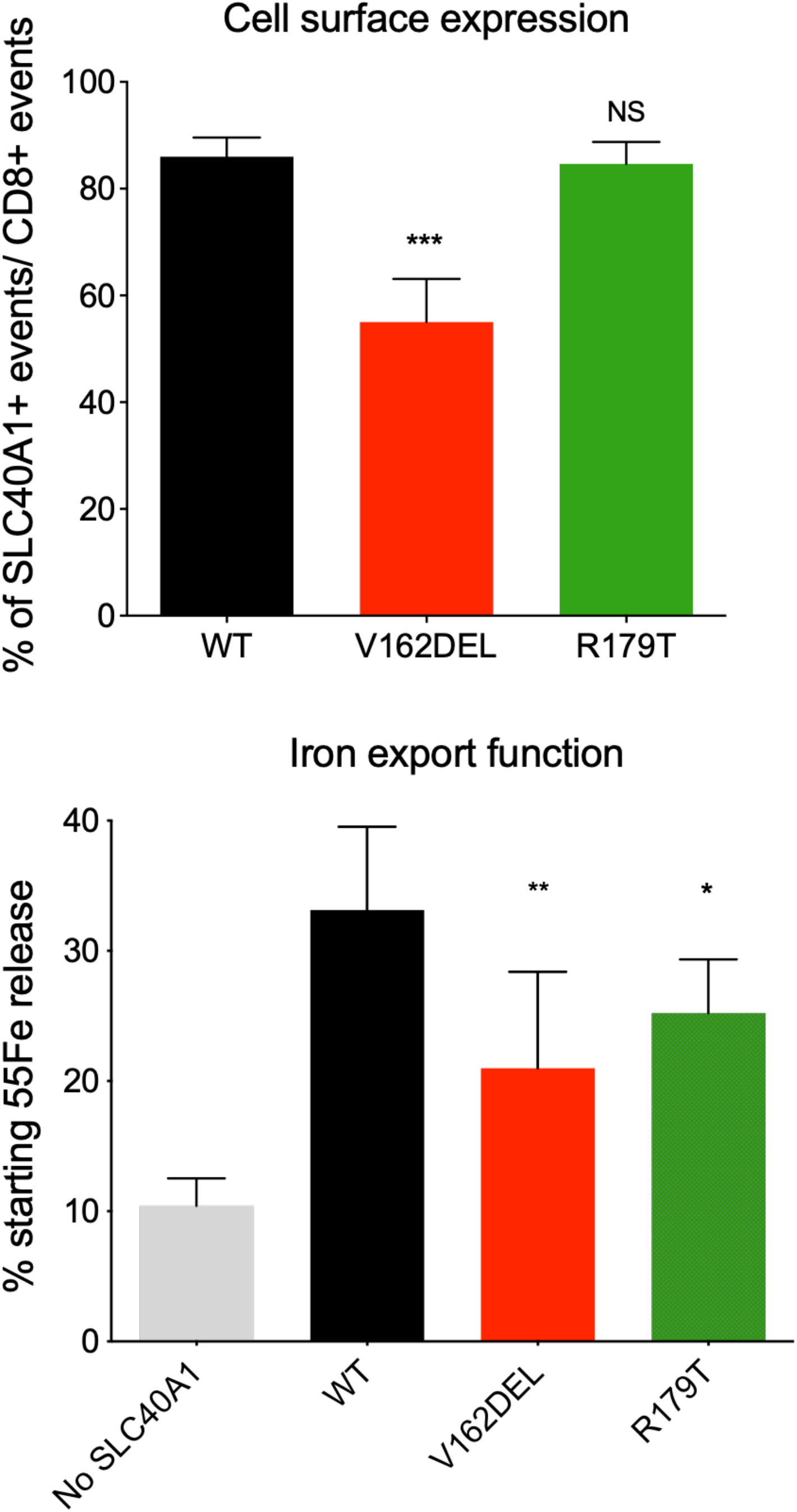
*In vitro* characterization of SLC40A1 missense mutations. (A) HEK293T cells were transiently transfected with the bicistronic pIRES2 plasmid encoding both full-length human FPN1-V5 and CD8. After 36 h, cells were double-stained for CD8 (APC) and the FPN1-V5 fusion protein (FITC, or PE) and analyzed by 2-color flow cytometry. Data are presented as percentages of FPN1-positive over CD8-positive events. (B) HEK293T cells were grown in 20 μg/mL ^55^Fe-transferrin for 24h before being washed and transiently transfected with wild-type or mutated SLC40A1-V5 expression plasmids. After 15 h, cells were washed and then serum-starved. The ^55^Fe exported into the supernatant was collected at 36h. Data are presented as percentage of cellular radioactivity at time zero. Each bar represents the means ± sd of 5 independent experiments. *P* values were calculated by a Student’s *t* test. ** p<0.01, *** p<0.001 and **** p<0.0001.

Still very close to the inner gate network, but deeper in TM5, we also identified Arg179. It introduces a positive charge in the dehydrated portion of the lipid bilayer, which can be considered as an unfavorable interaction, but this is compensated by the attraction of lipid head groups of the membrane toward the arginine residue. This attraction was observed at a significant rate, especially with POPC (frequency of contact of 30 % with POPC in the two simulations, while contact with POPE were observed with a frequency of 14 % and 5 %). This may actually cause localized bilayer deformation and the formation of an aqueous pathway, while ensuring thermodynamic stability, as already discussed in several articles (31, 32). In one of the two MD-POPC/POPE/CHOL simulations, a significant contact with Chol (frequency of 21 %) was also observed (Figure 1B).

### *In vitro* evaluation of five positively charged SLC40A1 amino acids that are likely to interact with glycerophospholipids

An alanine scanning mutagenesis was used to verify the functional importance of Lys90, Arg179, Arg365, Lys366 and Arg371, and consolidate the assumption of protein-lipid interfacial residues in close proximity to the human SLC40A1 inner gate. The mutants were tested for both plasma membrane expression and iron export. The single amino acid Val162del deletion, which is known to reduce cell surface localization of SLC40A1 (35), was used as negative control in all experiments.

A bicistronic construct was used to evaluate the concurrent plasma membrane expression of human FPN1 (conjugated with a V5 epitope) and cluster of differentiation 8 (CD8) in transiently transfected HEK293T cells on flow cytometry, as previously reported (13). An important difference was observed in the proportions of SLC40A1-WT^+^/CD8^+^ and SLC40A1-V162del^+^/CD8^+^ cells (p<0.001), confirming that the p.Val162del mutant causes SLC40A1 mislocalization. Comparable decreases were observed with the p.Lys90Ala, p.Arg365Ala, p.Lys366Ala and p.Arg371Ala mutants, whereas no differences were seen between cells transfected with the SLC40A1-WT/CD8 or SLC40A1-Arg179Ala/CD8 constructs (Figure 2A).

The *in vitro* activity of the five SLC40A1 mutants was assessed by measuring the release of radioactively labeled iron (^55^Fe) from transiently transfected HEK293T cells, following a well-established protocol (16, 33, 34). As shown in Figure 2B, HEK293T cells transiently transfected with a pcDNA3.1 plasmid encoding the wild type SLC40A1-V5 fusion protein displayed a 3-fold increase in iron release than cells transfected with the commercial pcDNA3.1-V5-His empty vector (No FPN1). None of the tested SLC40A1 mutants was not able to export ^55^Fe in amounts comparable with WT SLC40A1 (p<0.05-0.01); the largest effect being attributed to p.Arg179Ala.

These results show that the removal of a basic side chain at either of the TM1-90, TM5-179, TM8-365, TM8-366 or TM9-371 positions reduces iron export, in a way that is not necessarily related to a decrease in the cell surface expression of SLC40A1.

### *In vitro* evaluation of the p.Arg179Thr substitution, currently classified as a variant of unknown significance

We very recently examined the clinical significance of virtually all the missense variations reported in the *SLC40A1* gene in literature, taking into account all available phenotypic, familial and functional data (8). We classified the p.Arg179Thr substitution as a variant of unknown significance, as it has been reported in single patient (or Italian origin) without evidence of tissue iron overload (19). We thought it interesting to test its impact on SLC40A1 activity in transiently transfected HEK293T cells. As shown in Figure 3, the p.Arg179Thr mutant was found to have a profile similar to that of the p.Arg179Ala one; *i*.*e*. it reduces the ability of SLC40A1 to export iron without causing protein mislocalization.

## DISCUSSION

Although MFS transporters carry a wide variety of substrates and the molecular details of their transport mechanism may differ, a clear requirement is the formation and breakage of non-covalent bonds (salt bridges and H-bonds) that hold the N- and C-domains together. In SLC40A1, these bonds stabilize the OF conformation, awaiting cytoplasmic iron translocation. The complete mechanism of iron export requires a succession of conformational states that seems to be fine-tuned by the binding of iron, calcium and protons (14, 15, 18, 35, 36). Plasma membrane lipids certainly also regulate the stability and functional activity of SLC40A1, as it is now suggested for a growing number of MFS proteins (12), but no direct evidence has been reported yet.

MD simulations on the OF human SLC40A1 3D structure allowed us to identify five positively charged amino acids interacting with lipid head groups in a POPC bilayer and in a lipid bilayer including POPC, POPE and CHOL, chosen to mimic a standard plasma membrane (Figure 1). Four of them (TM3-Lys90, TM8-Arg365, TM8-Lys366, TM9-Arg371) are part of a cytoplasmic ring that surround the inner gate. Lys90 is even part of motif A, which is a hallmark of MFS superfamily and is known to form different gating interactions (11, 37). In human SLC40A1, motif A is located between TM2 and TM3 in the N-domain. It is critical for stability of the iron exporter facing the extracellular milieu (13–15). The fifth positively charged amino acid has a more distinctive positioning as it corresponds to a basic residue (Arg179 in TM5) which is more deeply embedded in the plasma membrane. This residue is adjacent to Arg178, which forms a salt bridge with Asp473 and actively participates to the charge/polar clusters that brings TM4, TM5, TM10 and TM11 in close proximity to create a stable inner gate (13– 15) (Figure 1).

The cellular effects of alanine-substituted mutants confirmed the functional importance of Lys90, Arg179, Arg365, Lys386 and Arg371 (Figure 2). The dynamic interaction of the four basic amino acids Lys90, Arg365, Lys366 and Arg371 with lipids could have an effect not only on the overall stability of SLC40A1 in the OF conformation but also on the formation/disruption of the critical salt-bridges in the vicinity of which they are located. The p.Arg179Ala mutant displayed different characteristics, as it was found to significantly reduce iron export without causing SLC40A1 mislocalization. This recapitulate some of our previous findings about mutants affecting amino acids (Asp157, Arg178, Arg489, Gln478, Asn174) involved in the formation of non-covalent interactions at the interface between the N- and C-domains (13, 16). Arg179 “snorkels” into the membrane, having its aliphatic chain in contact with the hydrophobic part of the lipid bilayer, while forming strong and persistent hydrogen bonds with lipid phosphates, resulting in a pulling of a patch of the lipid headgroup interface towards the hydrophobic core (Figure 1B, bottom). Consistent with other studies (31, 32), this whole phenomenon should reduce the energetic penalty associated with such a deep burial of an arginine in the bilayer. The position of Arg179, next to Arg178, raises the question of the potential influence of its dynamic interactions with phospholipids and the resulting local deformation of the bilayer on the conformational switch associated with the transport cycle. Moreover, worth noting is that Arg179 can also bind the oxygen atom of CHOL, which fits into a deep groove formed with TM1.

The interaction between charged residues and glycerophospholipids has been presented as a way for MFS transporters to refine the energetic barriers imposed by salt-bridges interactions. The amine headgroup of phosphatidylethanolamine (PE) has most specifically been described as a modulator of sugar and multidrug transporters (bacterial LacY, bacterial LmrP, bacterial XylE, human OAT1) (30, 38–41). Whatever the case, PE has been viewed to compete with gating residues, eventually in motif A, to facilitate discrete conformational changes. It is more generally admitted that MFS are highly sensitive to their lipid environment, which can modulate their stability, functional activity and (for some) oligomerization (12). Here, we have not addressed the issue of conformational changes that might be favored by the specific action of lipids on the inner gate salt-bridge network. We rather illustrate some indirect influence that lipids could play on this network, due to its proximity to positively charged residues whose interaction with the lipid polar heads may contribute to stabilize the protein in the outward open state. Nevertheless, we observed fluctuations in the Arg88-Asp157 salt-bridge in one of the two MD simulations performed in the POPC/POE/CHOL mixed lipid bilayer (but not in the simulations performed with POPC lipid molecules only) (Supplementary Data 2). Disruption of the Arg88-Asp157 salt-bridge may be associated with the specific interaction of the external Lys90 with POPE (Supplementary Data 3). On another hand, the particular position of Arg179, deep in the bilayer and close to the central cavity where iron sits during the MFS-related translocation mechanism, and its association with a local plasma membrane deformation could be a critical point for conformational switching.

As stated in the introduction, the p.Arg179Thr substitution has been reported in a 41-year-old Italian male with unexplained hyperferritinemia. Serum ferritin level at diagnostic was 819 µL, contrasting with normal transferrin saturation (39%). We previously classify this amino acid change as a variant of unknown significance, asking for further clinical, functional or structural data (8). We here demonstrate that it reduces the SLC40A1 iron export function, and thus corresponds to a new loss-of-function mutation. Its behavior is similar to that of the recurrent p.Arg178Gln mutation (16). This completes our previous findings (8, 13, 16, 34, 42, 43), pointing that the loss of a basic side chain in the cytosolic part of transmembrane helix 5, in the vicinity of the SLC40A1 inner gate and in contact with lipids, may also be responsible for FD.

In summary, this study highlights the existence of protein-glycerophospholipid interactions in the vicinity of two charged/polar clusters that play a critical role in the formation of the SLC40A1 inner gate. This can help ensure the thermodynamic stability of the OF conformation, and also prime the iron transporter for discrete conformational changes. It confirms that the molecular mechanisms of FD are not restricted to missense mutations decreasing SLC40A1 expression at the cell surface, and thus its iron export function, but also involve missense mutations affecting SLC40A1 activity within the plasma membrane, due to the loss of essential intra- or inter-domains non-covalent interactions, or interactions with surrounding lipids. Other studies are warranted to investigate in depth the dependence of FPN1 on lipids, and extend further our ability to interpret rare *SLC40A1* missense variants.

## Disclosure of Conflicts of Interest

The authors declare that they have no conflicts of interest.

## Data availability statement

Data available on request from the authors.

## Acknowlegments

This work was performed using High-Performance Computing (HPC) resources from Grand Equipement National de Calcul Intensif (GENCI) / Centre Informatique National de l’Enseignement Supérieur (CINES) (Grant 2023-A0150314556 on Adastra MI250 machine).

We thank Jean-Paul Mornon for insightful discussions.

## Notes

**Fundings:** This work was supported by grants from the “Institut Brestois Santé Agro Matière” (IBSAM - AAP2020; Modulofer), the Gaetan Saleun Association and the GR-Ex Laboratory of Excellence (reference ANR-11-LABX-0051). The GR-Ex label is funded by the IdEx “Investissements d’avenir” program of the French National Research Agency (reference ANR-18-IDEX-0001).

### Competing Interest Statement

The authors have declared no competing interest.

