## SupplementaryData for "Insights into the role of glycerophospholipids on the iron export function of SLC40A1 and the molecular mechanisms of ferroportin disease"

### Supplementary Data

**Supplementary Figure 1.** Root Mean Square Deviations (RMSD) of the protein C-alpha atoms were calculated along MD simulations trajectories using as reference the initial cryo-EM conformation (PDB: 6W4S). Different systems were considered unsing either a mixed POPC/POPE/CHOL bilayer (left panel) or a POPC bilayer (right panel). For each system, two replicas are represented in black and red.

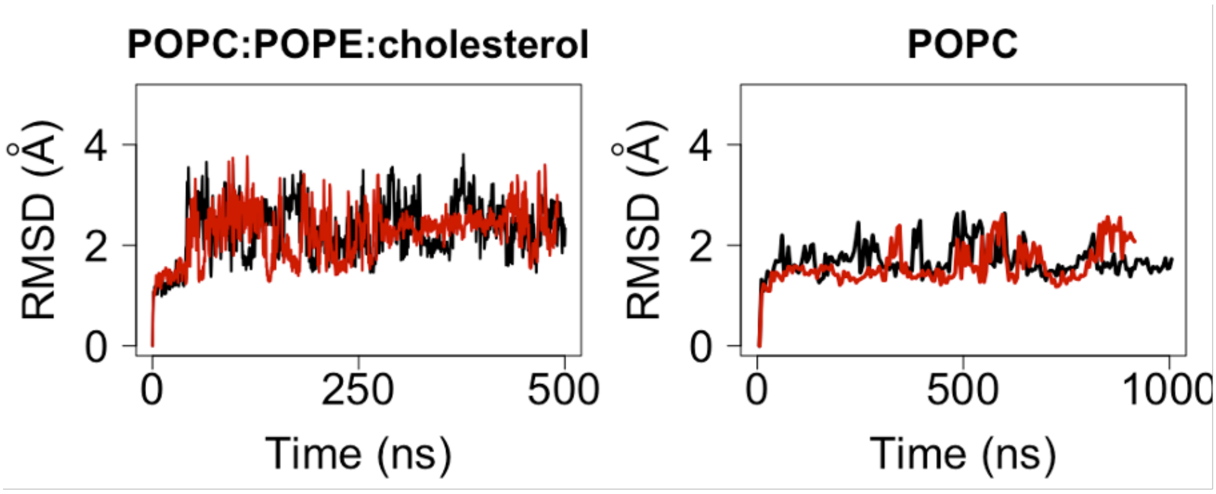

**Supplementary Figure 2.** Evolution, along the MD simulations, of the distances between atoms involved in noncovalent bonds. Distances were plotted (using R) as a function of the trajectories time for pairs of atoms in residues potentially involved in noncovalent bonds. The interaction reported here are based on the groups depicted in Fig. 1. The distance of 5 Å is considered as a reference threshold, below which bonds can be observed. For convenience, amino acids are described by a 1-letter code, instead of the 3-letter code used in the main text.

**POPC (MD1 and MD2)**

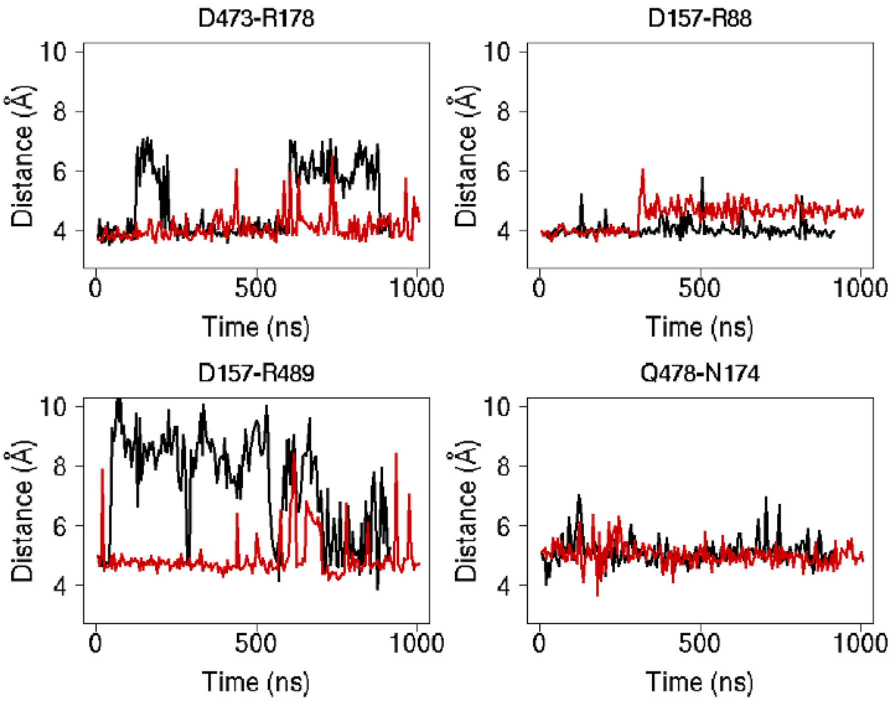


**POPC:POPE:Chol (MD3 and MD4)**

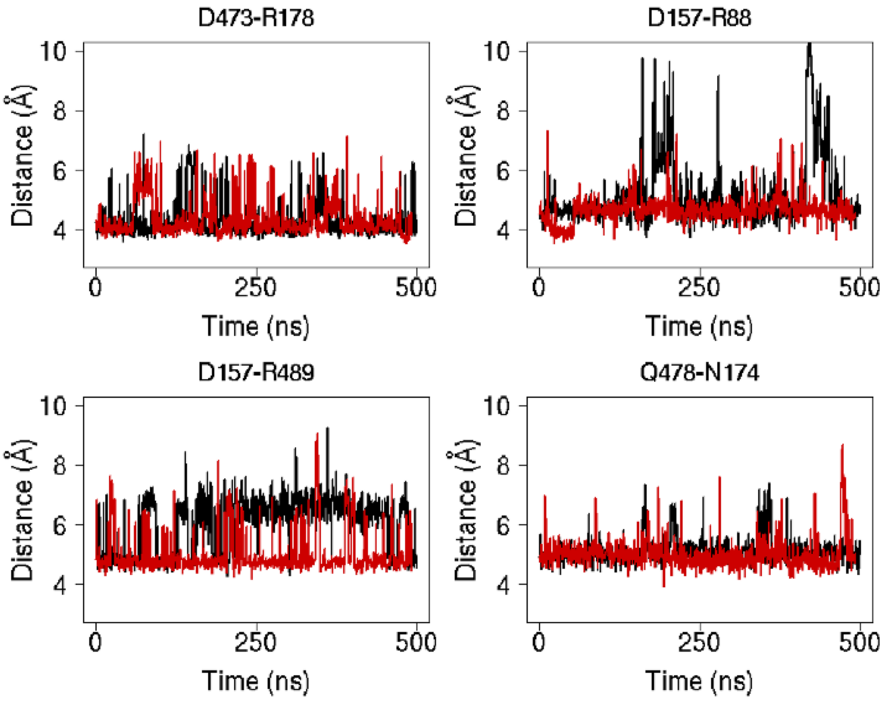

**Supplementary Figure 3.** Evolution of the R88-D157 interactions along MD simulations in the mixed POPC/POPE/CHOL bilayer, illustrated with two frames in which the salt-bridge between the two amino acids is present (50 ns) and absent (420 ns). The contacts of Lys90 with POPC and POPE are highlighted.

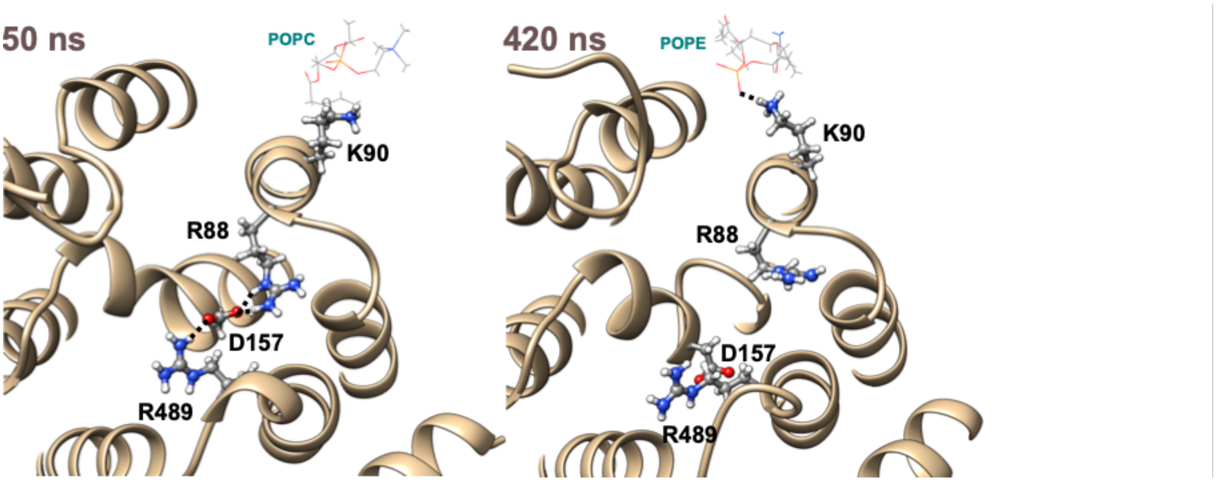
